# Deficit in observational learning in experimental epilepsy

**DOI:** 10.1101/2022.06.28.497932

**Authors:** Thomas Doublet, Antoine Ghestem, Christophe Bernard

**Affiliations:** INS - Institut de Neurosciences des Systèmes, Aix Marseille Université, 13005 Marseille, France

**Keywords:** observational learning, social behavior, spatial memory, animal model

## Abstract

Individuals use the observation of a conspecific to learn new behaviors and skills in many species. Whether observational learning is affected in epilepsy is not known. Using the pilocarpine rat model of epilepsy, we assessed learning by observation in a spatial task.

The task involves a naïve animal observing a demonstrator animal seeking a reward at a specific spatial location. After five observational sessions, the observer is allowed to explore the rewarded space and look for the reward.

Although control observer rats succeed in finding the reward when allowed to explore the rewarded space, epileptic animals fail. However, epileptic animals are able to successfully learn the location of the reward through their own experience after several trial sessions.

Thus, epileptic animals show a clear deficit in learning by observation. This result may be clinically relevant, in particular in children who strongly rely on observational learning.

## 1 INTRODUCTION

Observational learning (OL) - learning through observing the behavior of others - may be the most basic form of learning, central for survival for many species^1,2^. OL involves a sequence of four steps: attention, retention, reproduction and motivation^1,3,4^. Human infants essentially rely on OL, but adults also use OL, for example to learn new skills^5^. Many other species can adjust their behavior by observing conspecifics, including rodents^6,7,2^. Whether OL is affected in a pathological context is not known.

Patients with epilepsy commonly suffer from cognitive deficits^8^, a phenotype that can be recapitulated in various experimental rodent models of epilepsy^9^. Most studies focused on specific types of memories, including declarative, verbal and figural memory in patients^10,8^, and spatial/non-spatial or fear memories in experimental models^11^. Whether OL is affected in human or experimental epilepsy is not known.

Here, we used a behavioral task involving OL of a conspecific to assess the ability of epileptic rats (pilocarpine model) to learn a space or a task by observation^12^.

## 2 METHODS

### Animals

Animals were kept in a 12h LD light cycle and fed ad libitum. They were housed in environmentally enriched cages in a humidity and temperature-controlled environment. Twenty male Long Evans rats were included in the present study (3-7 months old at the time of testing, Charles Rivers Laboratories). All procedures took place during the light cycle.

Ten rats (213-244g) receiving intraperitoneal (i.p.) injections of pilocarpine hydrochloride (320mg/kg) 30 minutes after an i.p. injection of N-methyl-scopolamine (2mg/kg) developed SE (status epilepticus), which was stopped by diazepam (10mg/kg) after 120 minutes^13^. The rats did the behavior task 1 month after the SE.

All experiments were done in accordance with Aix-Marseille Université and Inserm Institutional Animal Care and Use Committee guidelines. The protocol was approved by the French Ministry of National Education, Superior Teaching, and Research, approval numbers APAFIS #30588- 2020121011005518v3 and #20325-2019041914138115v2.

### Experimental Design

We used an experimental design and protocol previously reported^12^. Briefly, experiments were conducted in a behavioral apparatus consisting of two square boxes: a transparent Plexiglas inner box within an opaque outer box. The space between the two boxes included 12 symmetrically distributed wells. An accessible but not visible reward (chocolate loops, Nestle) was placed in one of the wells.

### Behavioral Testing

All rats were familiarized with the experimental transparent inner box environment daily for at least three sessions of thirty minutes each (as shown in Figure 1). This allowed the inner box to be experienced directly, while the outer box could only be observed.

**FIGURE 1.**
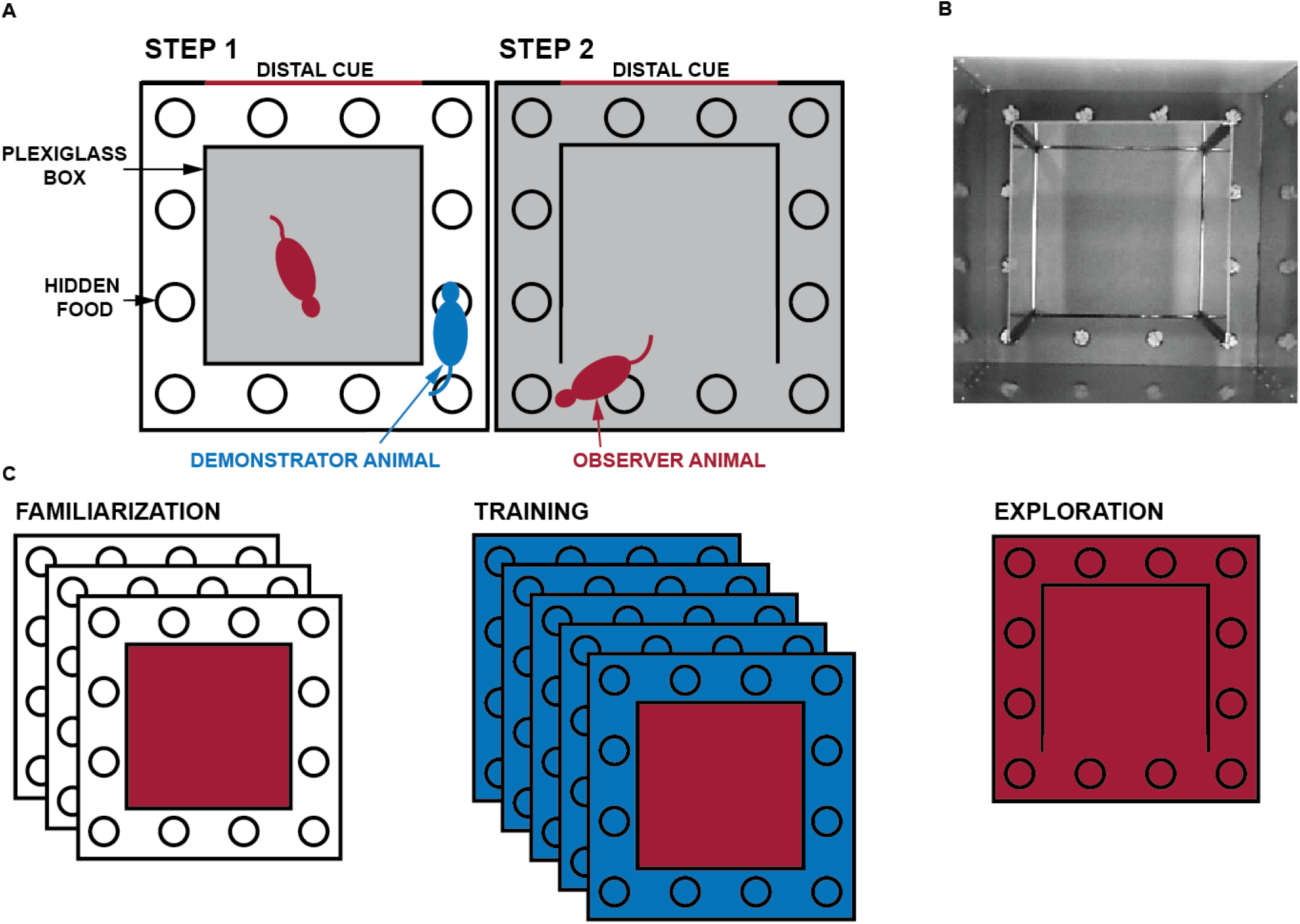
Experimental design. (A) The experimental environment consisted of a transparent inner box and an opaque outer box. The gray areas indicate the regions explored by the tested rat. In step 1, the observer (red) watches the demonstrator (blue), which has been trained to remember the location of a reward hidden in one of the 12 wells. (B) Image of the experimental apparatus with the lower wall of the transparent inner box open. The reward is covered with gravels. One of the four walls of the opaque outer box is white and provides a distal cue to the animals. (C) Schematic representation of the experiment. The familiarization phase, in which the experimental animal is confined to the inner box, is followed by the observational training phase, in which it can observe the demonstrator animal navigating the outer space. During direct exploration, the observer animal is allowed to navigate in the observed space. One session is held daily, for a total of 9 sessions (3 for familiarization, 5 for observational training, and 1 for direct exploration). The red and blue areas correspond respectively to the space that the observer and demonstrator animals can directly explore.

Subjects were divided into naive (n=5) and observer animals (n=15). Naive animals were tested for finding rewards without observational training. After at least twenty consecutive successful trials of finding the reward, the naive animals were considered as demonstrator animals. Each observer animal was then allowed to observe a demonstrator animal when the latter performs the task of seeking and getting the reward (Figure 1A-B). OL consisted of five daily demonstrations for five consecutive days with the observer in the inner box (Figure 1C). After OL completion, the observer rat was allowed to explore the outside space to seek and find the reward. The outside space was entered through the opening of a plexiglass wall opposite the reward well.

A performance was considered successful if the observer animal made no mistakes. A mistake was counted as active digging in an unrewarded well. A separate cohort of observers was tested without reward available to the animals during direct outside exploration.

### Data Analysis

All data were analyzed using the average time taken to find the reward from entering the outside space, the total number of mistakes made, and the percentage of successful animals for each trial. All values were expressed as mean ± standard error of the mean (SEM). All behavioral data were analyzed similarly to previously reported^12^. Effect sizes and confidence intervals (CI) are reported as: Effect size [CI width lower bound; upper bound].

Statistical significance was set at p < 0.05 p < 0.01 “*” and p < 0.001 “***”.

## 3 RESULTS

### 3.1 Observational learning in epileptic and control animals

We used the OL task already validated in control rats (Figure 1)^12^. Figure 2 shows the progression of success for a naive control animal in this task. A success is defined as finding the reward on the first try without foraging in other wells. The probability of finding the reward by chance is 8.3% (1 well out of 12). The probability of success on the first attempt for naive animals was comparable to chance (0%). The percentage of success increased in naive animals as the number of attempts increased: (attempt #1) 0% ± 0 (mean percentage ± SEM); (#2) 60% ± 24.5; (#3) 40% ± 24.5; (#4) 60% ± 24.5; (#5) 80% ± 20; (#6) 60% ± 24.5; (#7) 80% ± 20; (#8) 60% ± 24.5; (#9) 80% ± 20; (#10) 100%; (#11) 60% ± 24.5; (#12) 80% ± 20; (#13) 100%; (#14) 100% and (#15) 100% (n=5). The number of errors and the time taken by the animals to find the reward during the first attempt were 2.8 ± 0.7 and 3910 ± 1516 s, respectively, which is not statistically different from the previous results^12^ (the unpaired mean difference is 0.8 [95.0%CI – 0.538, 2.34] and 2.39e+03 [95.0%CI – 6.21e+02, 4.97e+03], respectively). Naive control rats learned the task until they reached a plateau.

**FIGURE 2.**
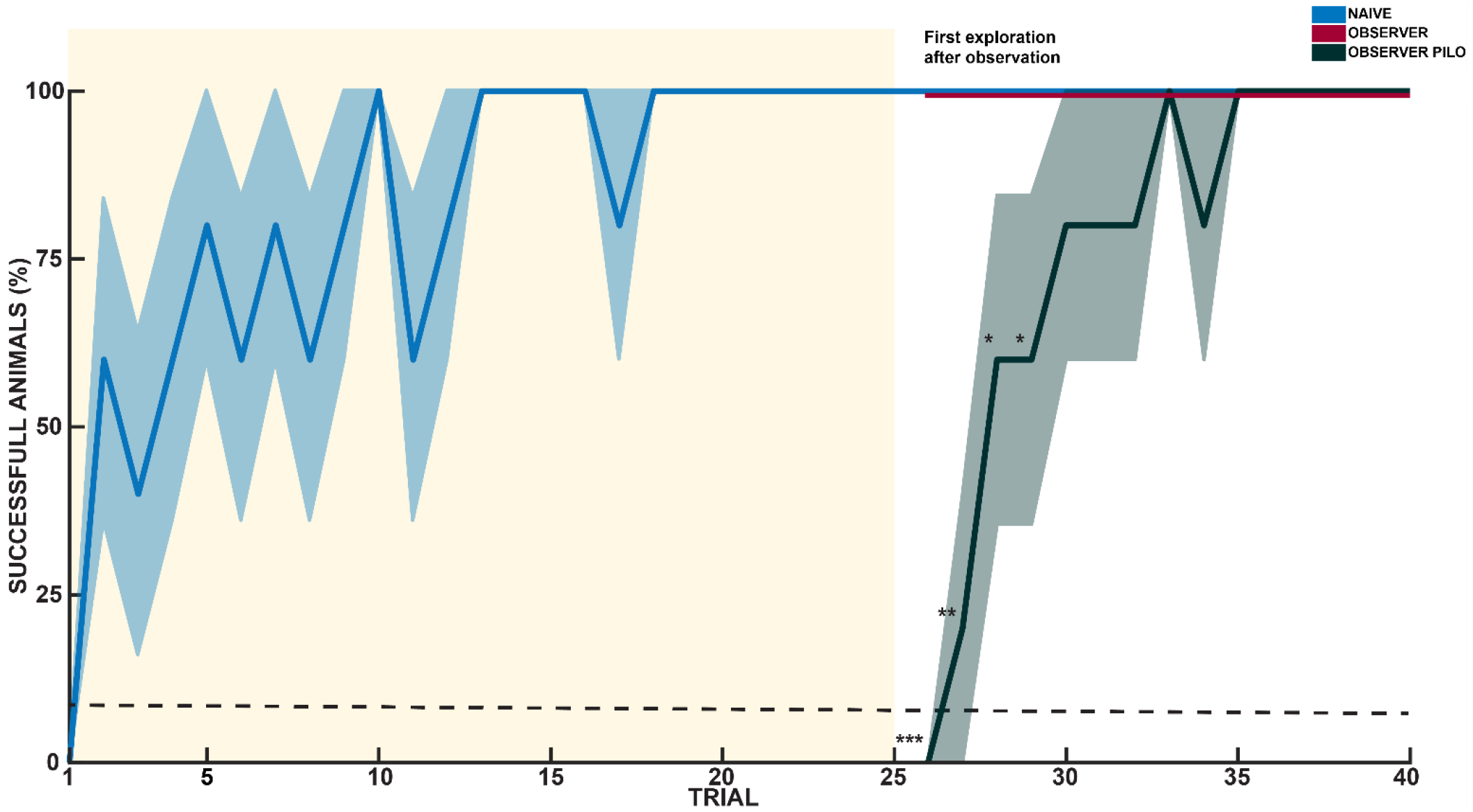
Epileptic animals fail to learn by observation. The percentage of animals successfully finding the reward is displayed as a function of trials. Naive control animals (blue, n=5) learn the location of the reward, and do not make any mistake after some time. They become demonstrators. After control observers (red, n=5) are first exposed to the rewarded space, they go straight to the reward; they have learnt by observation. Epileptic observers (black, n=5) fail to find the reward during the first exposure, but learn the task as naive control rats. Error bars are mean ± standard error of the mean (SEM). The black dashed line represents success by chance (8.3%). * p < 0.05, ** p < 0.01, *** p < 0.001.

To investigate whether control and epileptic animals can learn the location of a hidden reward by observation, observer rats watched demonstrators as they were running straight to the reward location (five trials daily for five consecutive days). Then, observers were allowed to directly explore the observed space to find the reward (see Figure 1).

In the control observer group, all animals successfully found the reward without error during their first direct exploration of the outside space (n=5) (Figure 2). This result is also not statistically different from what has been reported previously^12^ (unpaired mean difference: 7.14 [95.0%CI 0.0, 14.3]). All subsequent direct explorations were also 100% successful (n=15 trials, five animals).

In the epileptic observer group, no rat successfully found the reward without error during the first direct exploration of the outside space (n=5) (Figure 2). This result is statistically different from that of the control observer group (92.9 [99.9%CI −1e+02, – 71.4). The difference in percentage of success between control and epileptic observers persisted for the second, third and fourth attempts (the unpaired mean difference is −80.0 [99.0%CI −1e+02, −40.0], – 40.0 [95.0%CI −1e+02, −20.0] and −40.0 [95.0%CI −1e+02, −20.0], respectively). The number of errors for epileptic observers during the first exploration is statistically different from the control observer group (the unpaired mean difference is 3.2 [99.9%CI 2.0, 6.0]) and similarly in the second exploration (the unpaired mean – difference is 2.6 [99.9%CI 0.4, 7.0]). The time taken by epileptic observer rats to complete the first trial was 6124 ± 729 s, which is statistically different from the control observer group (1118 ± 747 s; the unpaired ‘ mean difference is 5.01e+03 [99.9%CI – 5.0e+02, 6.8e+03]).

Thus, unlike control observers, epileptic observers failed in finding the reward during the first exploration of the observed space.

### 3.2 Epileptic animals can learn the spatial memory task

Although epileptic animals failed in the observational learning task, they may have retained some information when observing the demonstrator, which would make them learn faster the location of the reward. Alternatively, they may have major cognitive impairment in spatial memory tasks, which would make them unable to learn the location of the reward.

Finding the reward by naive control rats requires seven trials before being consistently successfully done^12^. The percentage of animals finding the reward was comparable in naive control and epileptic observer rats during the seven first explorations. The time taken by naive control and observer epileptic rats to complete the first seven trials was also not statistically different. The number of errors in the first seven explorations was not statistically different between these two groups, except for the second trial (the unpaired mean difference is 2.0 [95.0%CI 0.2, 4.6]). In this case, the epileptic rats made more errors than the naive ones (2.6 ± 1.2 and 0.6 ± 0.4, respectively). However, after seven trials, epileptic observer animals were able to consistently find the reward as naive controls.

We conclude that epileptic animals can learn the location of the reward (spatial memory task) as well as non-observer control rats, and that observing a demonstrator does not make them learn the task faster.

### 3.3 Learning the location of the reward is independent of olfactory cues

To rule out reward localization by olfaction, the reward was removed after observational training before the first direct exploration in a second cohort of epileptic observer animals.

The difference between the two epilepsy groups was not significant. At the first exposure, the percentage of success was identical (0%), and the number of mistakes made was 3.8 ± 0.9 (n=5) in the unrewarded epileptic animals and 3.2 ± 0.7 (n=5) in the rewarded epileptic animals (the unpaired mean difference is 0.6 [95.0%CI −1.8, 2.4]). The time required to find the reward during the first exposure was also not significantly different, 4601 ± 1410 s and 6124 ± 729 s, respectively (unpaired mean difference is – 1.52e+03 [95.0%CI −4.62e+03, 9.78e+02]).

Rewarded and unrewarded epileptic observer animals showed similar performance, ruling out a possible olfactory influence on task success during the initial direct exploration.

### 4 DISCUSSION

The main finding of the study is that epileptic rats fail to learn by observation, although they succeed in the spatial memory task.

In humans, learning by observation requires, as a first step, to be attentive to the demonstrator^1,3,4^. Attentional deficits has been extensively described in epileptic patients and animal models^14,15,16,17^. Attentional deficits may explain failure in observational learning. However, we did not note a qualitative difference in the attention paid to the demonstrator between control and epileptic rats.

In this observational task, the hypothesis is that the observer builds a cognitive spatial map of the outside space. This map is used to navigate straight to the reward although the animal has never been directly exposed to this space. In experimental epilepsy, the hippocampus, which is critical for spatial memory, is characterized by numerous alterations^11,18^. Yet, epileptic animals can learn the spatial task used here when exposed to the new environment. This suggests that the formation of the cognitive spatial map by observation is very sensitive to such alterations, and/or that reward motivation helps the learning process when the animal is actively exploring the novel environment.

### 5 CONCLUSION

We provide the first evidence that observational learning in a spatial task is deficient in an experimental model of epilepsy. Given the importance of observational learning for humans throughout life, it will be important to test observational learning in patients with epilepsy, in particular in children.

## ACKNOWLEDGMENTS

Thomas Doublet gratefully acknowledges the support of the Ligue Francaise Contre l’Epilepsie – LFCE (Bourses LFCE 2020). The project received funding from Excellence Initiative of Aix-Marseille University – A*Midex, a French “Investissements d’Avenir” programme, and ANR-19-CE14-0036-01 “Stress”.

## ETHICAL PUBLICATION STATEMENT

We confirm that we have read the Journal’s position on issues involved in ethical publication and affirm that this report is consistent with those guidelines.

## AUTHOR CONTRIBUTION

**Thomas Doublet:** Conceptualization; Investigation (lead); formal analysis; funding acquisition; visualization, writing – original draft. **Antoine Ghestem**: Data acquisition. Investigation. **Christophe Bernard**: Supervision; resources; writing – review and editing.

